# Synergistic Activity of Repurposed Peptide Drug Glatiramer Acetate with Tobramycin Against Cystic Fibrosis *Pseudomonas aeruginosa*

**DOI:** 10.1101/2022.02.03.478806

**Authors:** Ronan A. Murphy, Matthew Coates, Sophia Thrane, Akshay Sabnis, James Harrison, Silke Schelenz, Andrew M. Edwards, Thomas Vorup-Jensen, Jane C. Davies

## Abstract

*Pseudomonas aeruginosa* is the most common pathogen infecting the lungs of people with cystic fibrosis (CF), causing both acute and chronic infections. Intrinsic and acquired antibiotic resistance, coupled with the physical barriers resulting from desiccated CF sputum, allow *P. aeruginosa* to colonise and persist in spite of antibiotic treatment. As well as the specific difficulties in eradicating *P. aeruginosa* from CF lungs, *P. aeruginosa* is also subject to the wider, global issue of antimicrobial resistance. Glatiramer acetate (GA) is a peptide drug, used in the treatment of multiple sclerosis (MS), which has been shown to have moderate anti-pseudomonal activity. Other antimicrobial peptides (AMPs) have been shown to be antibiotic resistance breakers; potentiating the activities of antibiotics when given in combination restoring and/or enhancing antibiotic efficacy. Growth, viability, minimum inhibitory concentration (MIC)-determination and synergy analysis showed that GA improved the efficacy of TOB against reference strains of *P. aeruginosa*, reducing TOB MICs and synergising with the aminoglycoside. This was also the case for clinical strains from people with CF. GA significantly reduced the concentration of TOB required to inhibit 50% (MIC_50_) of viable cells (from 1.69 [95%CI 0.26-8.97] to 0.62 [95%CI 0.15-3.94] mg/L, p=0.002) and inhibit 90% (MIC_90_) (from 7.00 [95%CI 1.18-26.50] to 2.20 [95%CI 0.99-15.03] mg/L, p=0.001) compared with TOB-only. Investigating mechanisms of GA activity showed that GA resulted in significant disruption of outer membranes, depolarisation of cytoplasmic membranes and permeabilisation of *P. aeruginosa* and was the only agent tested (including cationic AMPs) to significantly affect all three.

## INTRODUCTION

*Pseudomonas aeruginosa* is a Gram-negative, rod-shaped bacterium found ubiquitously in the environment and frequently associated with opportunistic infections (burns, wounds, eye-infections). *P. aeruginosa* is the most common infecting bacteria in the lungs of people with cystic fibrosis (CF) (1). Cystic fibrosis is a genetic, life-limiting disorder and, while a multi-system illness, lung disease causes the majority of morbidity and mortality in people with CF; impaired mucociliary clearance leads to chronic bacterial infection, significant inflammation and bronchiectasis resulting, ultimately, in respiratory failure (2–4). Forty-one percent of adult CF patients (>16 years) in the UK are chronically infected with *P. aeruginosa*, with a peak of 54% in the 36-39 age cohort (1). *P. aeruginosa* is of particular concern due its ability to evade antibiotic treatments (via both innate and acquired mechanisms) leading to its designation as an ESKAPE pathogen by the World Health Organisation (5–7).

The aminoglycoside, tobramycin is one of the most commonly used anti-pseudomonal antibiotics in cystic fibrosis. Administration is either intravenous (IV) or inhaled directly to the airway (1). Limitations of the IV route include a narrow therapeutic index requiring concentration monitoring to avoid oto- and nephrotoxicity. The inhaled route allows high local levels with less systemic exposure, but once infection is chronic, drug efficacy is modest and eradication rare, a limitation shared with other agents at this disease stage. In part, this may relate to heterogeneity of deposition within partially obstructed airways, giving rise to subtherapeutic drug concentrations. Few new treatments are being developed for chronic *P. aeruginosa*, and the growing adult population means this unmet need will persist into the era of novel modulator drugs targeting the underlying cellular defect in CF (8,9).

A potential solution to the need for novel *P. aeruginosa* therapy, among the dearth of new antibacterial treatments, is the deployment of antimicrobial peptides (AMPs), small biological molecules, usually consisting of 10-50 amino acid residues, which form part of innate immune systems produced widely across all kingdoms of life (10–12). They frequently function via interactions with the bacterial membrane which lead to cell wall weakening, thinning and/or permeabilisation and rapid cell death (10,13–15). The Gram-negative membrane consists of an outer membrane, containing outwards protruding lipopolysaccharide, and a cytoplasmic membrane with a thin peptidoglycan layer between. AMPs are of interest not only because of their own potential antimicrobial activity but also their possible use as antibiotic adjuvants; given in combination with antibiotics, they may reduce concentrations of the latter required for antimicrobial activity (16). This property, often referred to as ‘antibiotic resistance breaking’, has been reported against *P. aeruginosa*: the human-derived AMP LL-37 and its derivatives with azithromycin, vancomycin (17,18), novel synthetic peptides with tobramycin and colistin (19–21). However, despite promising results, many AMPs have significant barriers to clinical use, including concerns around cytotoxic effects (13,22,23).

Glatiramer acetate (GA) is an immunomodulator drug currently in clinical use in the treatment of multiple sclerosis (24–26). It is produced by the random polymerisation of four amino acids (L-glutamate, L-lysine, L-alanine, and L-tyrosine residues) and therefore structurally resembles AMPs. We have shown previously that GA has moderate antimicrobial activity comparable to LL-37 against *P. aeruginosa* – both reference strains and clinical strains from people with CF, optimal activity occurring at 50mg/L (15,27). GA has other properties in common with naturally occurring antibiotic resistance breaking AMPs being cationic, <150 amino acid residues long and exerting antimicrobial effects over an extremely short time course to a level similar to that of the AMP LL-37 (26,28).

Investigating existing drugs which are not conventional antibiotics for their potential use as antimicrobials, known as ‘repurposing’, is a growing area of interest and research (29,30). As well as addressing the paucity of new antibiotics being developed, repurposing of existing drugs has several other advantages: approved drugs have historical safety data and repurposing may minimise time and costs associated with deployment to the clinic due the availability of previous trials and research (30).

As with other members of the aminoglycoside family, tobramycin targets the bacterial protein synthesis machinery and thus requires uptake into the bacterial cytoplasm for activity (31). Given its widespread use in CF, preserving, restoring and/or enhancing the efficacy of tobramycin is of particular interest and importance. As it has also been previously subject to ‘breaking’ by antimicrobial peptides, we examined tobramycin in combination with GA, to assess the peptide as a potential antibiotic resistance breaker in cystic fibrosis strains of *P. aeruginosa*. We quantified the impact of GA on bacterial survival at a range of tobramycin concentrations and assessed its direct effects on outer and cytoplasmic membranes to elucidate mechanisms of action.

## MATERIALS AND METHODS

### *Pseudomonas aeruginosa* Isolates

Reference *P. aeruginosa* strains PAO1, PA14 and PAK and 11 clinical *P. aeruginosa* isolates were used in this study. Clinical strains were selected from the CF Bacterial Repository at the National Heart and Lung Institute, Imperial College London which consists of bacteria isolated from airway samples of people with cystic fibrosis at the Royal Brompton Hospital, London (Table 1). Bacteria were stored in Microbank vials (Pro-Lab Diagnostics, UK) at -80°C. Clinical strains were selected with a range of antibiograms according to clinical lab results using EUCAST clinical breakpoints. Isolates were revived onto Cetrimide agar (Merck, UK) for confirmation of purity and grown overnight at 37°C followed by subculture onto LB agar (Merck, UK). Single colonies were inoculated into Mueller-Hinton broth (Merck, UK) and incubated overnight at 37°C with agitation at 200r.p.m.

**Table 1.** Clinical *P. aeruginosa* from people with cystic fibrosis isolated at Royal Brompton Hospital, London used in this study.

| Strain | Sample Type | Mucoidy | Multidrug Resistance | Tobramycin | Ceftazidime | Ciprofloxacin | Colistin | Meropenem |
| --- | --- | --- | --- | --- | --- | --- | --- | --- |
| RBH550 | Spontaneous Sputum |  |  | Sensitive | Sensitive | Sensitive | Sensitive | Sensitive |
| RBH461 | Cough Swab |  | MDR | Resistant | Resistant | Resistant | Sensitive | Resistant |
| RBH490 | Spontaneous Sputum |  | MDR | Resistant | Resistant | Resistant | Sensitive | Resistant |
| RBH519 | Spontaneous Sputum | Mucoid |  | Resistant | Sensitive | Resistant | Sensitive | Sensitive |
| RBH072 | Spontaneous Sputum |  |  | Resistant | Sensitive | Resistant | Sensitive | Sensitive |
| RBH294 | Spontaneous Sputum | Mucoid |  | Sensitive | Sensitive | Sensitive | Sensitive | Sensitive |
| RBH899 | Spontaneous Sputum |  |  | Sensitive | Sensitive | Sensitive | Sensitive | Sensitive |
| RBH750 | Spontaneous Sputum | Mucoid |  | Sensitive | Sensitive | Sensitive | Sensitive | Sensitive |
| RBH065 | Spontaneous Sputum |  |  | Resistant | Sensitive | Resistant | Sensitive | Sensitive |
| RBH422 | Spontaneous Sputum |  |  | Sensitive | Sensitive | Resistant | Resistant | Sensitive |
| RBH982 | Spontaneous Sputum |  |  | Sensitive | Sensitive | Sensitive | Sensitive | Sensitive |

### Bacterial Growth Curves and Colony Counts

Late-exponential phase cultures of *P. aeruginosa* were harvested by centrifugation (3,500g for 15mins) and resuspended in fresh Mueller-Hinton broth. Tobramycin solutions and bacterial cultures were prepared in Mueller-Hinton broth at double the desired concentrations and then mixed with cultures 1:1. Starting bacterial optical density at 600nm (OD_600_) was 0.05. Tobramycin concentrations were tested stepwise in Log2 increments. Tobramycin-bacteria solutions were vortexed briefly and aliquoted in triplicate 200μL volumes into a sterile 96-well plate (ThermoScientific, UK). Glatiramer acetate (Biocon, India) was added to the remainder of the tobramycin-bacterial suspension to a final GA concentration of 50mg/L, the concentration previously identified as optimal (15). Samples were vortexed and aliquoted to 3 further wells of the 96-well plate, also in 200μL volumes. Untreated bacteria, bacteria treated with 50mg/L GA only and uninoculated Mueller-Hinton wells were also included. Plates were sealed with optically clear, breathable seals (4titude, UK) and incubated in a FLUOstar Omega platereader (BMG Labtech, UK). Optical density OD_600_ was measured every 30mins, overnight at 37°C with shaking at 200r.p.m.

After overnight growth, triplicate wells were pooled, centrifuged, supernatant removed and pellets resuspended in sterile phosphate buffered saline (PBS) (Merck, UK). Serial 1:10 dilutions were performed in sterile PBS and diluted cultures were spotted onto LB agar plates using the Miles-Misra method; 20μL volumes with ≥10 technical repeats per condition were plated (32). Plates were allowed to dry at room temperature and placed in 37°C incubator overnight. Spots with between 2 and 20 individual colonies were quantified.

### Outer Bacterial Membrane Disruption

Disruption of the outer bacterial membrane of *P. aeruginosa* was measured using the fluorescent probe 1-N-Phenylnaphthylamine (NPN) (Merck, UK) (33). Late-exponential phase cultures of *P. aeruginosa* were washed twice and adjusted to an OD_600_ of 0.5 in 5mM HEPES buffer. In the wells of a black microtitre plate (ThermoScientific, UK), adjusted cultures were combined with NPN (final concentration 10μM) along with either no treatment (buffer only), 50mg/L GA, 2mg/L colistin (CST), 4mg/L tobramycin (TOB) or 16mg/L LL-37 (Merck, UK), to a final volume of 200μL. Agent concentrations were chosen based on EUCAST breakpoint concentrations (CST and TOB) or previously published active concentrations (GA and LL-37, (34) and (35) respectively). Each experiment consisted of triplicate wells of each condition. Fluorescence was measured in a platereader at excitation 355nm, emission 460nm every 30secs for 15mins. NPN Uptake Factor was calculated as ([Fluorescence of Sample with NPN – Fluorescence of Sample without NPN] / [Fluorescence of buffer with NPN – Fluorescence of buffer without NPN]).

### Cytoplasmic Membrane Depolarisation

Depolarisation of the cytoplasmic membranes of *P. aeruginosa* isolates was measured using the fluorescent probe 3,3’-Dipropylthiadicarbocyanine Iodide (DiSC_3_(5)) (Thermo Scientific, UK) (36). Late-exponential phase cultures of *P. aeruginosa* were washed twice and adjusted to an OD_600_ of 0.05 in 5mM HEPES-20mM glucose. DiSC_3_(5) was added to the bacterial cultures to a concentration of 1μM and aliquoted to the wells of a black microtitre plate. The fluorescent signal of the dye was allowed to quench for 30mins in the dark followed by addition of equal volumes of buffer (No Treatment), GA (for 50mg/L), CST (2mg/L), TOB (4mg/L) or LL-37 (16mg/L). Each experiment consisted of triplicate wells of each condition. Fluorescence was measured in a platereader at excitation 544nm, emission 620nm every 30secs for 1hr.

### Bacterial Membrane Permeabilisation

Permeabilisation of the bacterial membranes of *P. aeruginosa* was measured using the fluorescent dye propidium iodide (PI) as per manufacturer’s instructions (Merck, UK). Late-exponential phase cultures of *P. aeruginosa* were washed twice and adjusted to an OD_600_ of 0.5 in PBS. Propidium iodide was added to the bacterial cultures to a concentration of 1μg/mL and treatments added: no treatment, 50mg/L GA, 2mg/L CST, 4mg/L TOB or 16mg/L LL-37. Cell were aliquoted to the wells of a black microtitre plate in 200 μL volumes. Each experiment consisted of triplicate wells of each condition. Fluorescence was measured in a platereader at excitation 544nm, emission 610nm every 30secs for 1hr.

### Statistical Analyses

Bacterial growth curves were plotted by transformation of OD_600_ data using natural log, of biological replicates, and displayed as median and 95% confidence intervals. OD_600_ at 18hrs and Area Under the Curve of biological replicates were compared by Kruskal-Wallis with Dunn’s multiple comparison (to untreated) and adjusted p values reported. Where the Kruskal-Wallis results were significant (p<0.05), individual TOB concentrations were compared, with GA versus without GA, via Mann-Whitney t test.

Synergy between the activities of glatiramer acetate and tobramycin was tested for each strain from colony counts, as previously described (37). Briefly, the Inhibition Effects in CFU/mL caused by 50mg/L GA (*E*_*A*_) and each TOB concentration tested (*E*_*B*_) were used to calculate Expected Effect (*E*_*Exp*_) of combining both treatments, where *E*_*Exp*_ = *E*_*A*_ + *E*_*B*_ – (*E*_*A*_ * *E*_*B*_). The Observed Effect (*E*_*OBs*_) of the combined treatments was compared to the calculated *E*_*Exp*_ whereby *E*_*OBs*_ > *E*_*Exp*_ indicates Synergy, *E*_*OBs*_ = *E*_*Exp*_ indicates Additivity/Independence and *E*_*OBs*_ < *E*_*Exp*_ indicates Antagonism. We report Synergy when the above criteria were met and the resulting values had non-overlapping 95% confidence intervals.

Inhibition of each strain by TOB-only and GA/TOB was calculated for each tobramycin concentration tested, as a percentage of the untreated cultures, using CFU/mL results. Inhibition curves were generated for each *P. aeruginosa* strain using Nonlinear Fit ([inhibitor] vs. response (three parameters)). Tobramycin concentrations required to inhibit 50% (MIC_50_) and 90% (MIC_90_) of viable bacteria were interpolated from the curves generated for TOB-only and GA/TOB. Median MIC_50_ and MIC_90_ for all clinical strains were compared using Wilcoxon t tests.

Effects of different treatments on reference strains were compared for median NPN Uptake Factor over 15mins, median DiSC_3_(5) release over 15mins and median Area Under the Curve of PI fluorescence intensity over 1hr by Kruskal-Wallis with Dunn’s multiple comparison (to untreated) and adjusted p values reported. Clinical isolates were compared using the same membrane metrics for No Treatment vs 50mg/L GA with Wilcoxon signed rank tests. All analyses were performed in GraphPad Prism (9.2). p<0.05 was considered significant.

## RESULTS

### Combining glatiramer acetate and tobramycin alters growth and viability of *Pseudomonas aeruginosa*

We anticipated that if GA possessed antibiotic potentiation properties, this would likely be demonstrable only with certain concentrations of TOB; in the presence of an effective concentration (> strain-specific MIC) of antibiotic, any impact of GA could not be visualized whereas presumably too low a concentration of antibiotic could limit the impact of a compound capable of potentiation. We first tested this with a relatively high-throughput approach using growth curves generated from serial OD_600_ measurements during overnight culture of PAO1, PA14 and PAK exposed to a range of tobramycin concentrations (0.06-4mg/L in log2 increments) with and without 50mg/L GA. No strain was inhibited by GA alone and the concentration of TOB alone which impacted growth differed across strains, PA01 being the most susceptible and PAK the least (Figure 1 & S1). Confirming our hypothesis, for each strain we could identify TOB concentrations at which GA potentiated its activity.

**Figure 1.**
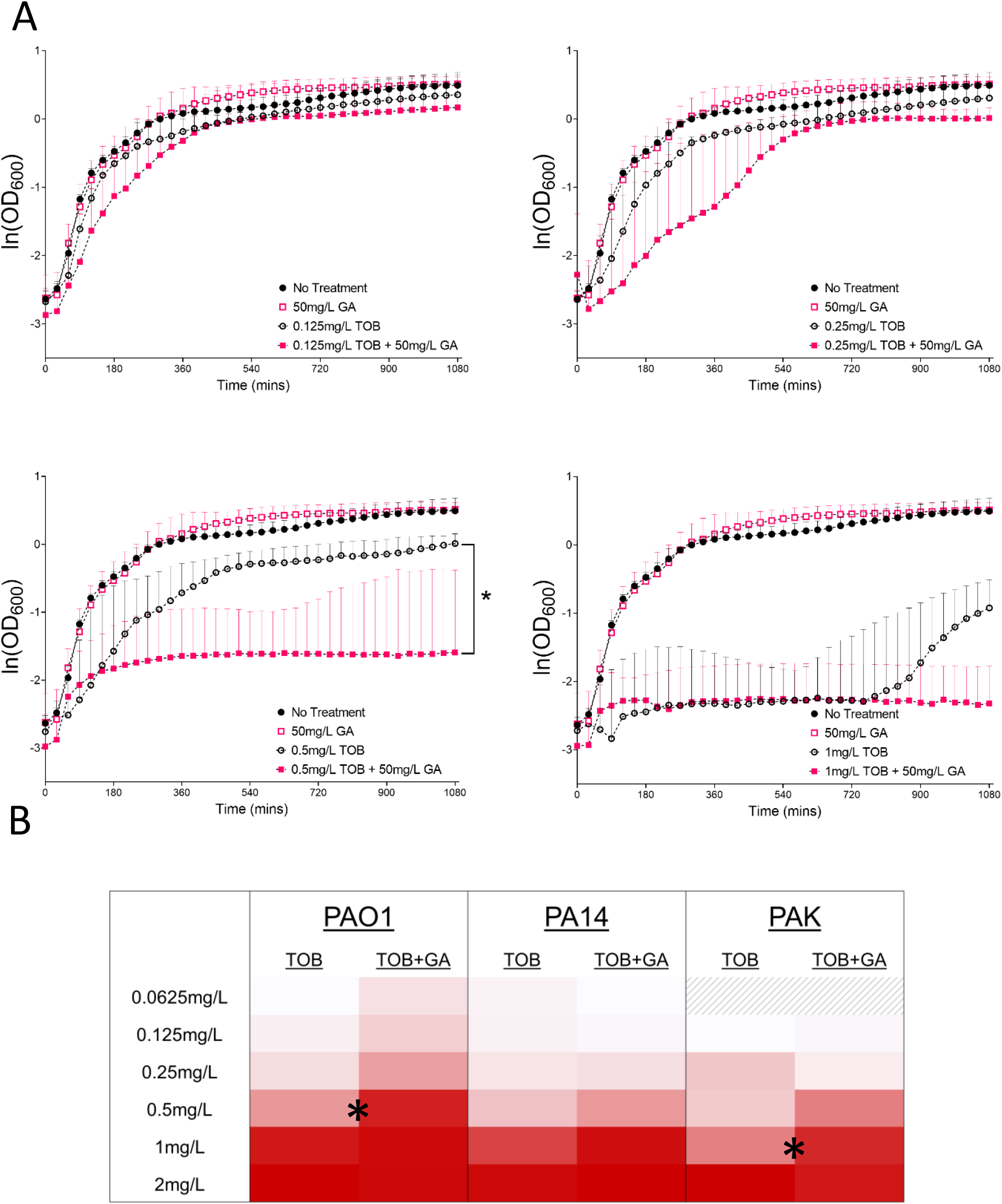
A. Semilog plots of overnight growth of *P. aeruginosa* PAO1 (natural logs of OD_600_, median and 95% confidence intervals) at increasing concentrations of TOB, with and without GA. Combination effect of GA/TOB was noted at 0.5mg/L TOB where the final OD_600_ of the growth curves were significantly lower for GA/TOB than TOB (p<0.05). B. Heat Map of Areas Under the Curve of PAO1, PA14 and PAK OD_600_ overnight growth curves at all TOB concentrations tested (median of at least three biological replicates). Darkness of red indicates lower AUCs and less bacterial growth. For PAO1 and PAK, GA/TOB resulted in significantly lower AUCs than TOB only at 0.5 and 1mg/L, respectively (p<0.05).

For strain PAO1, this concentration was 0.5mg/L TOB (Figure 1A). Median OD_600_ at 18hrs of PAO1 cultures treated with GA/TOB (0.5mg/L) was significantly lower than that of cultures treated with this concentration of TOB alone (0.21 [95%CI 0.17-0.78] vs 1.11 [95%CI 0.74-1.35]). p<0.05). Area Under The Curve analyses confirmed this significant effect (p<0.05; Figure 1B). For strain PAK, AUC analysis demonstrated potentiation of TOB by GA at 1mg/L (p<0.05) (Figure 1B, S1). While similar analyses for PA14 did not reach statistical significance, a visual trends suggestive of potentiation was observed, also at 1mg/L TOB (Figure S1).

Next, we sought to confirm and extend the observations from growth curve assays by colony counting. Viable colony counts were performed on overnight cultures after treatment with GA, TOB or GA/TOB with at least three biological replicates at each antibiotic concentration. GA alone did not affect CFUs of any strain and consistent with the findings above, CFUs were reduced more by GA/TOB co-treatment than by TOB alone (Figure 2). Of note, significant impact was observed at a greater range of TOB concentrations than had been observed with growth curve screening.

**Figure 2.**
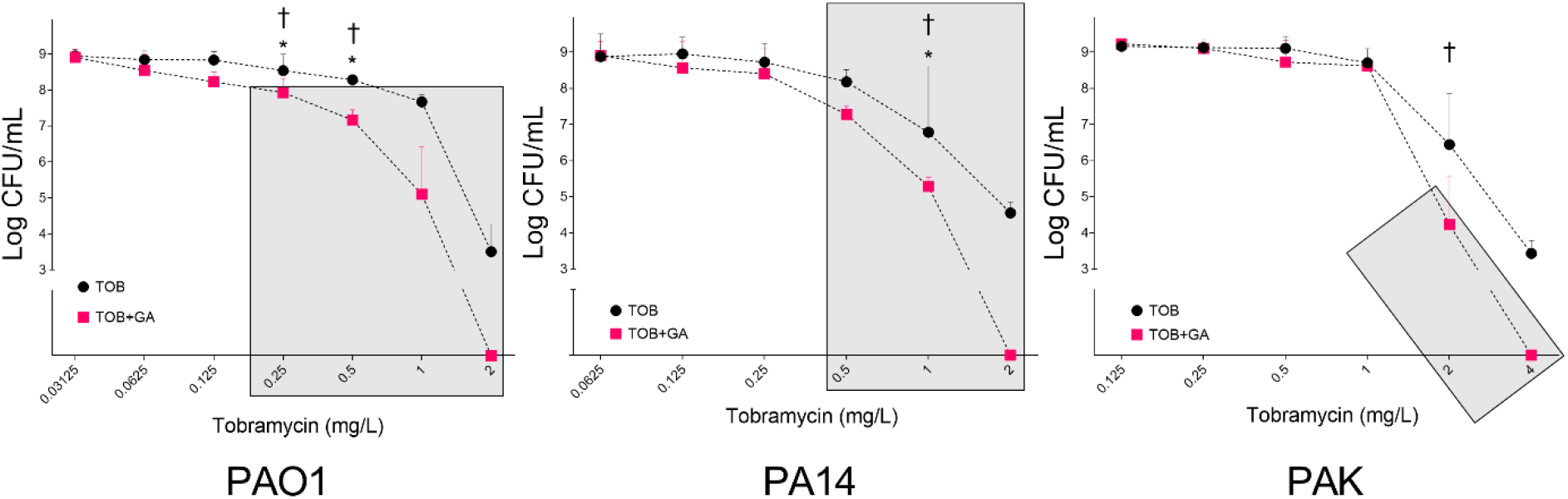
Colony forming units of strains PAO1, PA14 and PAK after overnight exposure to TOB and GA/TOB at increasing TOB concentrations (median with 95% confidence intervals). * indicates concentrations for which GA/TOB resulted in significantly fewer viable bacteria that TOB (p<0.05). † indicates the TOB concentrations where GA/TOB displayed synergy (*E*_*OBs*_ > *E*_*Exp*_). Shaded areas indicate results which where CFU/mL were significantly lower than untreated *P. aeruginosa* (data not shown on graphs: stable CFU/ml value as seen with lowest concentrations of TOB) for a given strain (p<0.05). For each strain, the highest GA/TOB concentration tested eliminated viable bacteria, whereas the same TOB-only treatment did not.

For PA01, combining GA with either 0.25 or 0.5mg/L TOB resulted in significantly fewer colonies than the same concentration of TOB-only (8.54*10^7^ [95%CI 1.38*10^7^-2.10*10^8^] vs 3.48*10^8^ [95%CI 1.41*10^8^-1.00*10^9^] and 1.49*10^7^ [95%CI 1.66*10^6^-2.85*10^7^] vs 1.94*10^8^ [95%CI1.33*10^8^-2.65*10^8^] respectively; p<0.05 for both). For both of these GA/TOB combinations, numerically fewer viable bacteria were recovered than with the 2-log higher TOB concentration administered alone (Figure 2). Furthermore, these TOB concentrations used alone did not significantly reduce CFU/ml compared with untreated cultures, whereas both GA/TOB concentrations had highly significant impacts (p<0.01) (Figure 2). For PA14, significant impact of the addition of GA was seen at 1 mg/L with >1log10 fewer CFU/ml (1.94*10^5^ [95%CI 2,050-3.50*10^5^] vs 6.10*10^6^ [95%CI 4.80*10^6^-4.19*10^8^]; p<0.05) (Figure 2). The impact on PAK was visually similar, but did not reach statistical significance (Figure 2).

We further analysed these CFU data to assess whether any impact of GA on TOB was additive or synergistic, the latter defined as the combined effect of the two agents being greater than would be predicted by the individual effects seen for each (*E*_*OBs*_ > *E*_*Exp*_). For PAO1 the combination of GA with both 0.25mg/L and 0.5mg/L TOB was synergistic; for PA14, synergy was observed for GA with 1mg/L TOB; for PAK GA with 2mg/L TOB was synergistic (Figure 2).

### Glatiramer acetate reduces the minimum inhibitory concentrations of tobramycin against reference strains of *P. aeruginosa*

For each of the three reference strains, colony count results were used to calculate the inhibition of *P. aeruginosa* (as a percentage of untreated cultures) caused by TOB and GA/TOB, at each TOB concentration tested, and inhibition curves generated (Figure 3). From the resulting fitted curves the minimum inhibitory concentrations of tobramycin required to reduce the CFU/mL by 50% (MIC_50_) and by 90% (MIC_90_) were interpolated when GA was present and absent (Figure S2).

**Figure 3.**
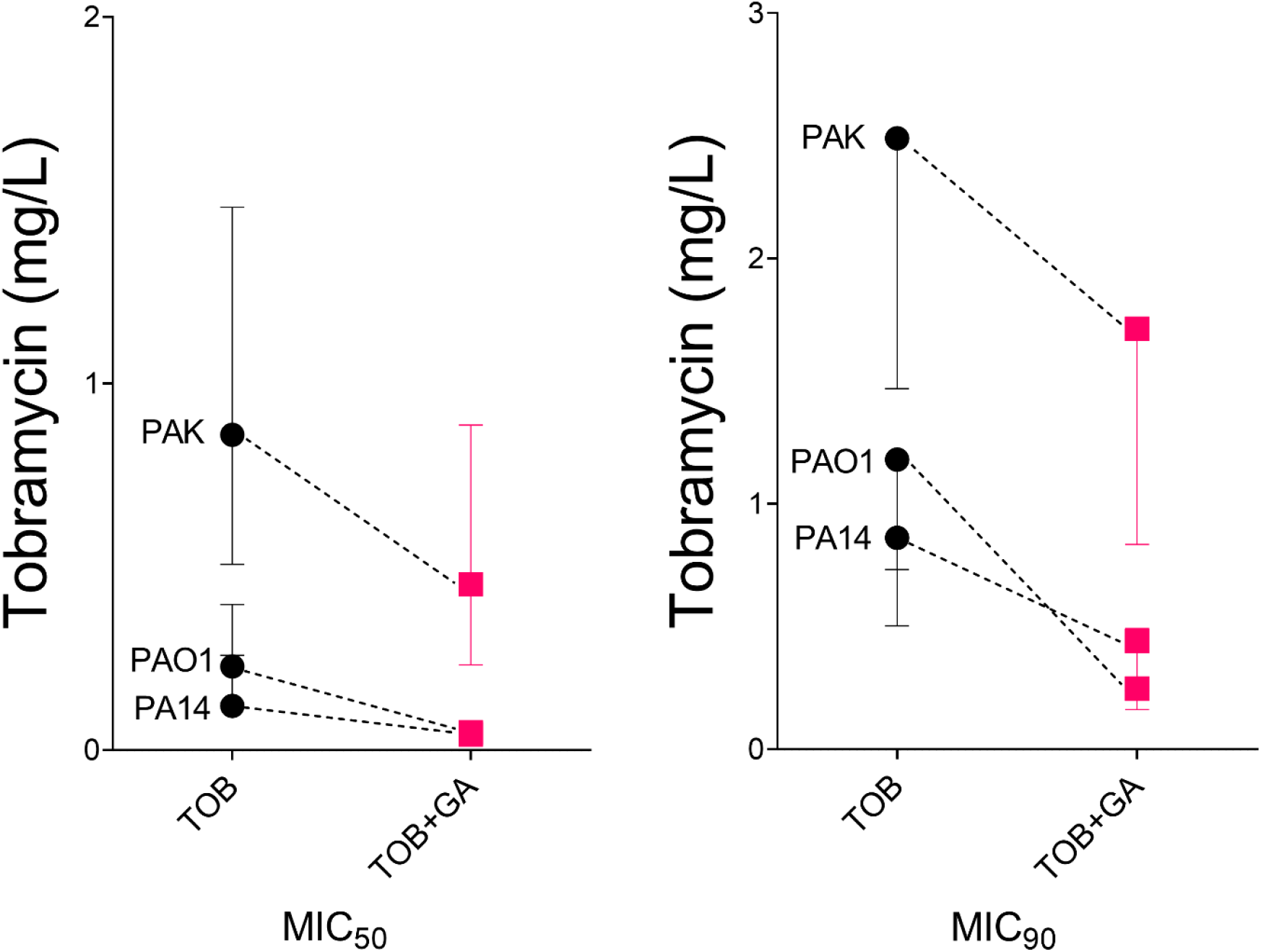
MIC_50_ and MIC_90_ values of TOB for strains PAO1, PA14 and PAK in the absence and presence of GA. Interpolated MIC values and their 95% confidence intervals are shown (from non-linear fit of inhibition curves of CFU/mL results derived using results from at least 3 biological replicates).

Fold reductions in MIC_50_ of 4.7, 2.8 and 1.9 and in MIC_90_ of 4.7, 1.9 and 1.5 for PA01, PA14 and PAK respectively were observed (Figure 3).

### Glatiramer acetate reduces the MIC values of tobramycin for clinical cystic fibrosis strains of *P. aeruginosa*

Eleven clinical isolates of *P. aeruginosa* from people with cystic fibrosis were selected (Table 1). As with the reference strains, clinical isolates were screened using growth curves, colony counts performed and MIC values interpolated from Inhibition Curves for each isolate (at least three biological replicates were performed). The sensitivity results generated for tobramycin via this method were consistent with the disc diffusion results reported by the clinical lab, using the EUCAST MIC breakpoint of 4mg/L for tobramycin.

Across the whole panel of 11 CF clinical *P. aeruginosa* tested here, GA significantly reduced both the MIC_50_ (median from 1.69 [95%CI 0.26-8.97] to 0.62 [95%CI 0.15-3.94] mg/L; 2.7-fold reduction, p=0.002) and MIC_90_ (median from 7.00 [95%CI 1.18-26.50] to 2.20 [95%CI 0.99-15.03] mg/L; 1.7-fold reduction, p=0.001) concentrations of tobramycin (Figure 4). Synergy analyses based on these results indicated that for 6 of the 11 clinical strains tested, there was at least one TOB concentration where the antibiotic synergised with GA, with the remainder having at least one TOB concentration for which GA was additive. Importantly, no antagonism was seen for any isolate at any concentration of TOB.

**Figure 4.**
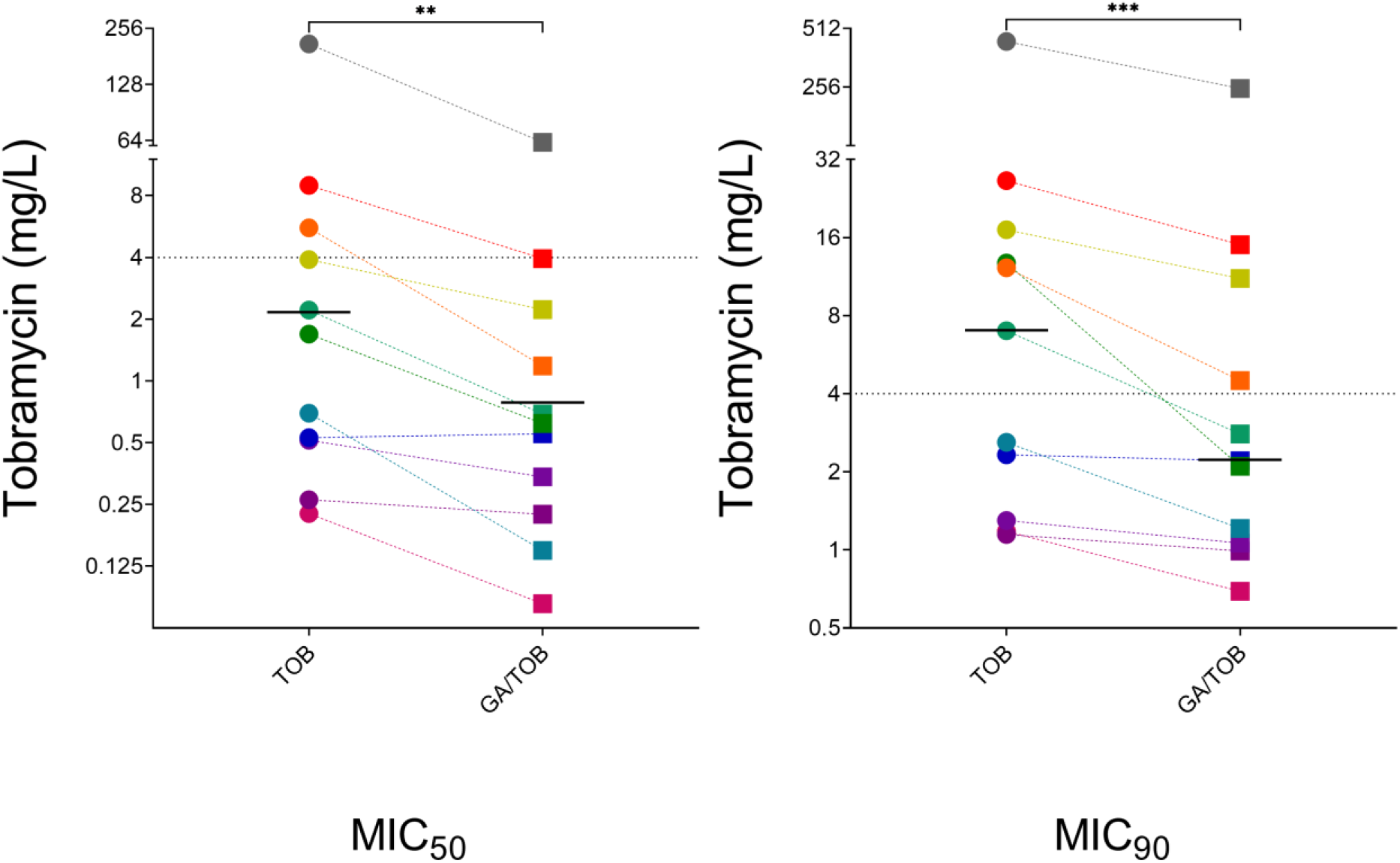
MIC_50_ and MIC_90_ results for TOB for 11 clinical *P. aeruginosa* strains from cystic fibrosis in the absence and presence of GA. Values were interpolated from non-linear fit of inhibition curves of CFU/mL results from at least 3 biological replicates for each strain. Across the 11 strains tested both MIC_50_ and MIC_90_ values of TOB were significantly reduced by co-administration of GA (p<0.01).

The 6 TOB-sensitive *P. aeruginosa* strains had an MIC_50_ for TOB alone of 0.52 [95%CI 0.23-2.21] mg/L which in the presence of GA was 0.28 [95%CI 0.08-0.69] mg/L (ns). Similarly for these strains, the MIC_90_ was 1.81 [95%CI 1.15-7.00] mg/L for TOB alone and 1.14 [95%CI 0.69-2.80 mg/L for GA/TOB (p<0.05). GA was not effective in reducing tobramycin MICs for every sensitive strain tested; one strain, RBH294, showed similar results for TOB effectiveness whether GA was present or not.

The 5 tobramycin resistant clinical *P. aeruginosa* isolates also demonstrated reductions in effective concentrations of tobramycin when GA was co-administered. The tobramycin MIC_50_ of this group was reduced by GA from 5.57 [95%CI 1.69-211.19] to 2.23 [95%CI 0.62-62.50] mg/L and MIC_90_ fell from 17.2 [95%CI 12.23-438.32] to 11.11 [95%CI 2.10-251.49] mg/L (neither of which reached statistical significance for this subgroup). GA reduced the tobramycin MIC_50_ by 2.72 [95%CI 1.75-4.70]-fold and the MIC_90_ 1.76 [95%CI 1.54-6.08] fold. For 4 of the 5 tobramycin-resistant strains tested, co-treatment with GA resulted in their MIC_50_ being decreased to <4mg/L, the EUCAST breakpoint concentration of tobramycin, indicating these strains would be considered tobramycin-sensitive in the presence of GA.

### Glatiramer acetate disrupts the outer bacterial membrane of *P. aeruginosa*

We next wished to explore the mechanism by which GA was potentiating the activity of TOB. The fluorescent probe, NPN, was used to measure disruption of the outer membrane of *P. aeruginosa* strains. Increased fluorescence indicates NPN is binding to the inner hydrophobic elements of the outer membrane, which are only accessible in disrupted membranes. Treatment with GA significantly increased median uptake of NPN by PAO1, PA14 and PAK, over 15mins. GA resulted in a 3.43 [95%CI 3.28-3.49]-fold increase in NPN uptake by reference strains of *P. aeruginosa*, compared with untreated cultures (p<0.0001) (Figure 5). Comparison with other antimicrobials showed that uptake of NPN from GA treatment was comparable with treatment with colistin (2mg/L) (3.99 [95%CI 3.71-4.79]-fold increase) and the membrane disrupting peptide LL-37 (16mg/L) (3.70 [95%CI 2.74-3.80]-fold increase), while treatment with the aminoglycoside antibiotic tobramycin resulted in 2.14 [95%CI 1.97-2.55]-fold increase in NPN Uptake Factor, compared to untreated *P. aeruginosa* cells (Figure 5).

**Figure 5.**
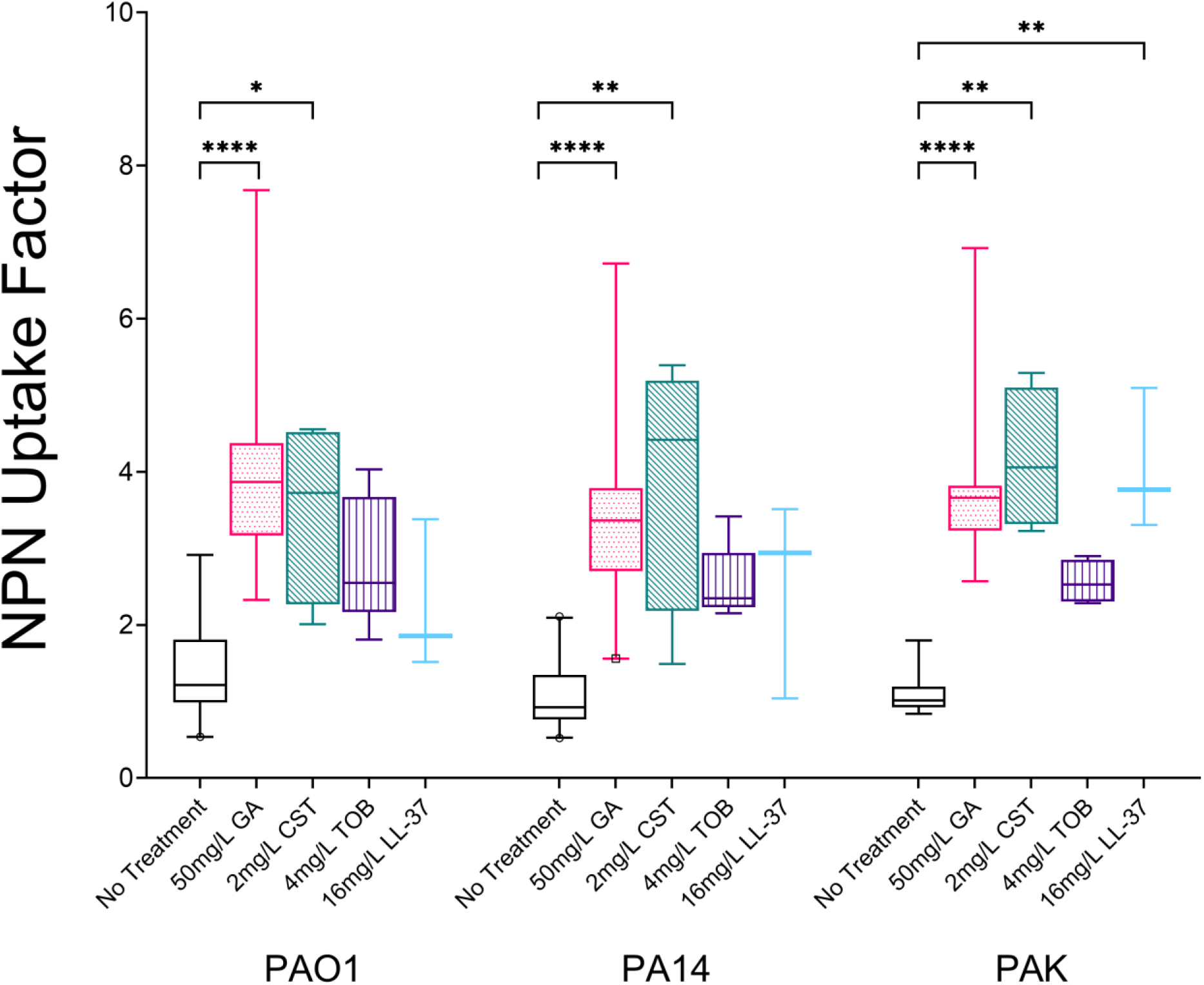
Comparison of uptake of NPN over 15mins by the outer membranes of strains PAO1, PA14 and PAK resulting from exposure to 50mg/L GA, 2mg/L CST, 4mg/L TOB and 16mg/L LL-37 (Box and Whisker plot, line at median with 5-95 percentiles. At least three biological replicates of each treatment were performed for each strain). GA caused a significant increase in outer membrane disruption for all three reference strains (p<0.0001).

Glatiramer acetate caused a significant increase in the uptake of NPN in the outer membranes for the group of 11 clinical *P. aeruginosa* strains from CF (median untreated 1.02 [95%CI 0.52-1.93] to GA-treated 2.45 [95%CI 1.38-4.02] (p=0.001) (Figure 6A).

**Figure 6.**
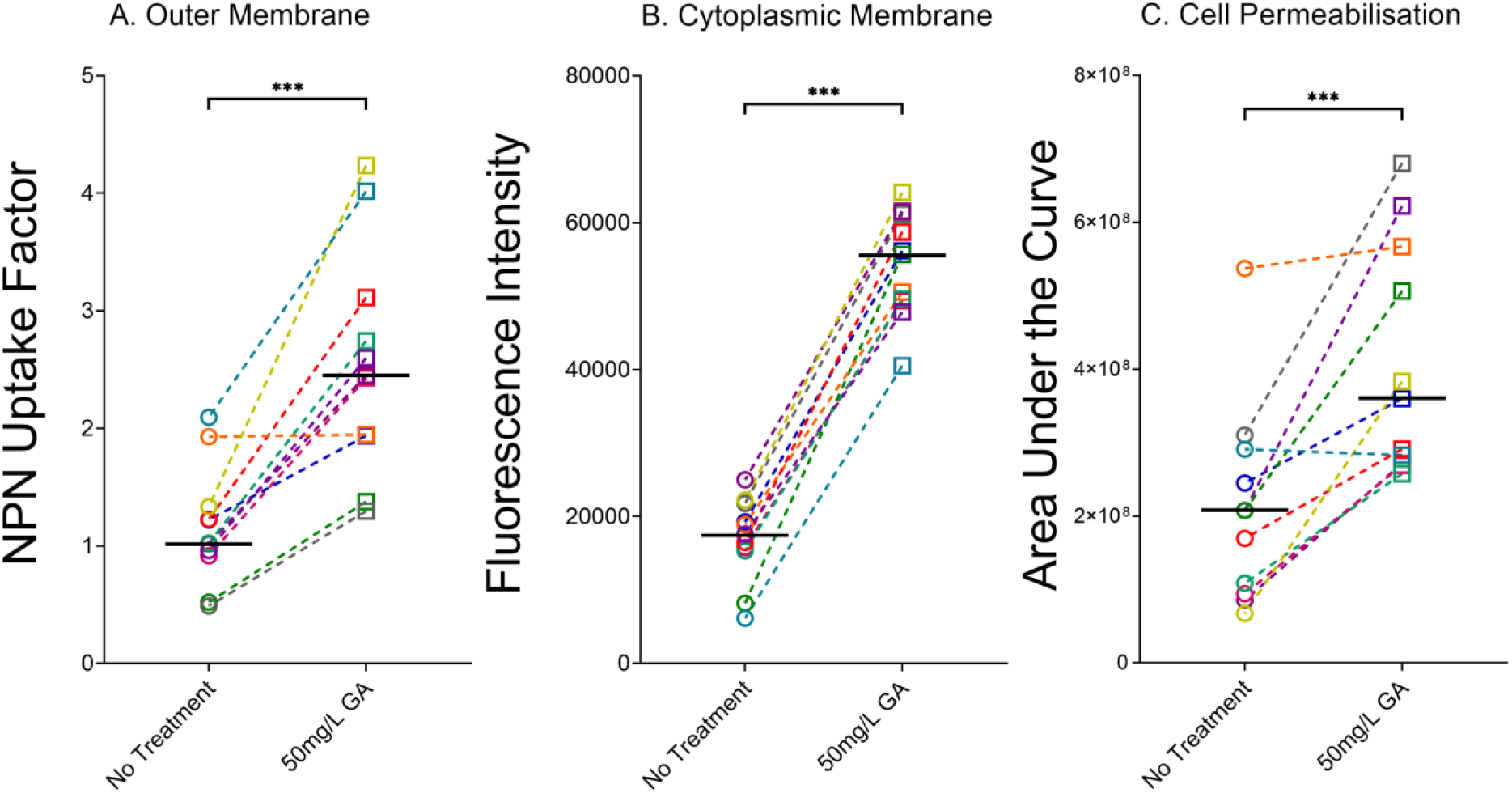
Effect of 50mg/L GA on the A. Outer Membranes B. Cytoplasmic Membranes C. Permeability of 11 cystic fibrosis *P. aeruginosa* strains. Each point represents a strain and is the median of at least three independent experiments. Line at median of all clinical strains tested. GA caused significant disruption of bacterial outer membrane, significant depolarisation of the cytoplasmic membrane and significant permeabilisation of the cell wall, compared to untreated bacteria (p<0.001).

### Glatiramer acetate depolarises the cytoplasmic membrane of *P. aeruginosa*

The fluorescent dye DiSC_3_(5) was used to measure depolarisation of the cytoplasmic membrane of *P. aeruginosa*. On application, the dye integrates into the intact cytoplasmic membrane; any subsequent intervention causing membrane depolarisation will lead to dye release and increased fluorescent signal. Treatment with GA resulted in rapid release of DiSC_3_(5) by PAO1, PA14 and PAK with a higher fluorescent signal produced than by TOB or CST. Examining the median fluorescence over the first 15mins of treatment, showed that GA resulted in each of the three *P. aeruginosa* reference strains releasing significantly more DiSC_3_(5) than the untreated cultures (p<0.0001) while, of the other agents tested, only LL-37 (16mg/L) caused significant cytoplasmic membrane depolarisation of PA14 and PAK (Figure 7).

**Figure 7.**
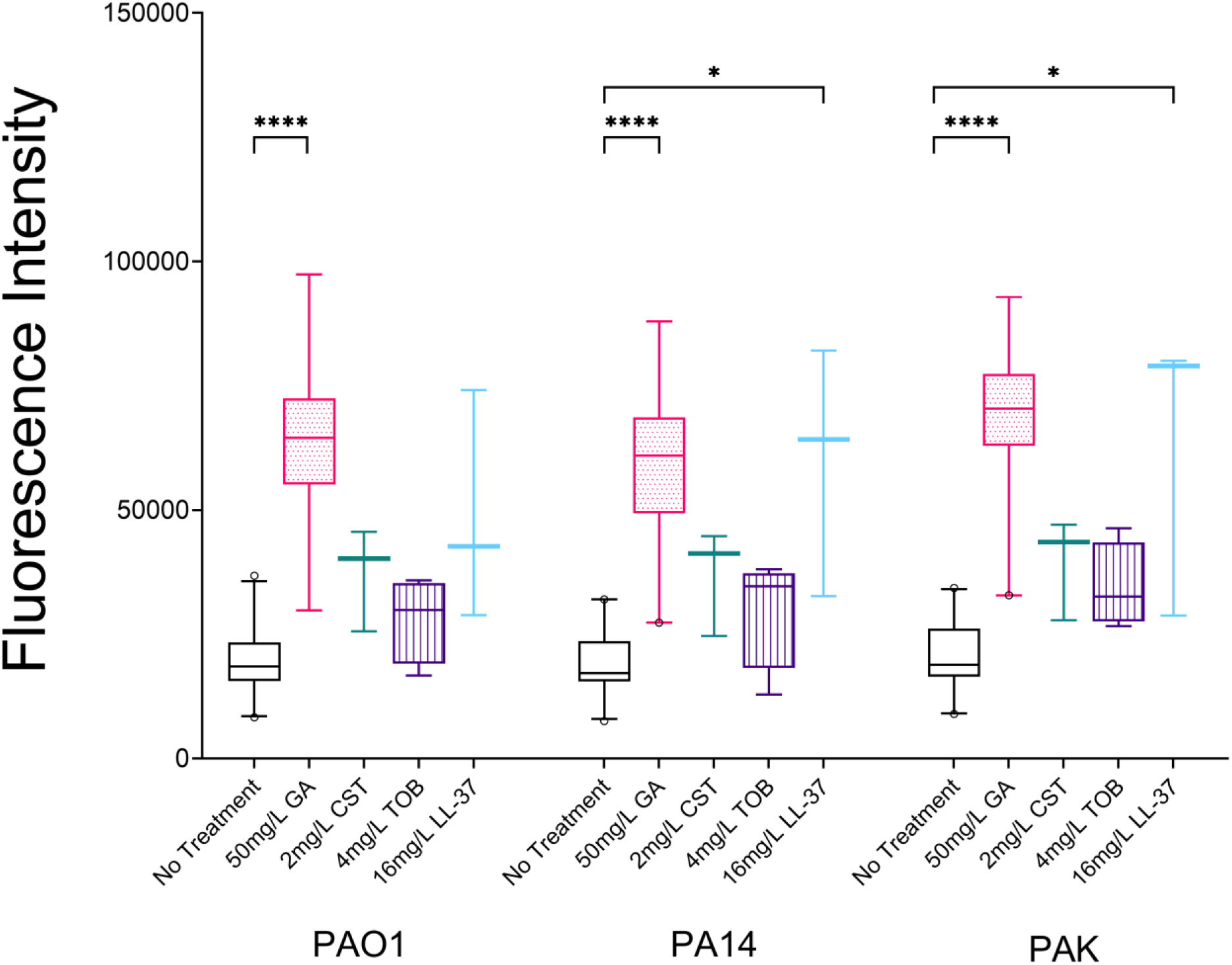
Comparison of release of DiSC_3_(5) over 15mins by the cytoplasmic membranes of strains PAO1, PA14 and PAK resulting from exposure to 50mg/L GA, 2mg/L CST, 4mg/L TOB and 16mg/L LL-37 (Box and Whisker plot, line at median with 5-95 percentiles. At least three biological replicates of each treatment were performed for each strain). GA caused a significant depolarisation of cytoplasmic membranes for all three reference strains (p<0.0001).

A similar effect was seen for the 11 clinical CF *P. aeruginosa* strains. Median DiSC_3_(5) release over 15mins was 3.19 [95%CI 2.68-6.32]-times higher with 50mg/L GA (p=0.001) compared with no treatment. Treatment with GA resulted in a minimum increase in dye release of 2.65 [95%CI 2.13-2.96]-fold while the highest level seen for any clinical strain was a 6.81 [95%CI 2.75-7.06]-fold increase over untreated (Figure 6B).

### Glatiramer acetate permeabilises *P. aeruginosa* cells

Having noted the ability of GA to perturb both the outer and cytoplasmic membranes of *P. aeruginosa*, the DNA-binding membrane impermeable dye, propidium iodide was used to test whether the bacteria were permeabilised by GA. Each *P. aeruginosa* reference strain was significantly permeabilised by 50mg/L GA compared to untreated cultures, assessed by AUCs of 1hr PI fluorescence (p<0.001) (Figure 8). None of the other agents resulted in permeabilisation with the exception of LL-37 (PA14 (p<0.001) and PAK (p<0.05)) (Figure 8). For the clinical strains GA resulted in a 2.37 [95%CI 1.05-3.18]-fold increase in permeabilisation (p=0.002 vs untreated cells) (Figure 6C).

**Figure 8.**
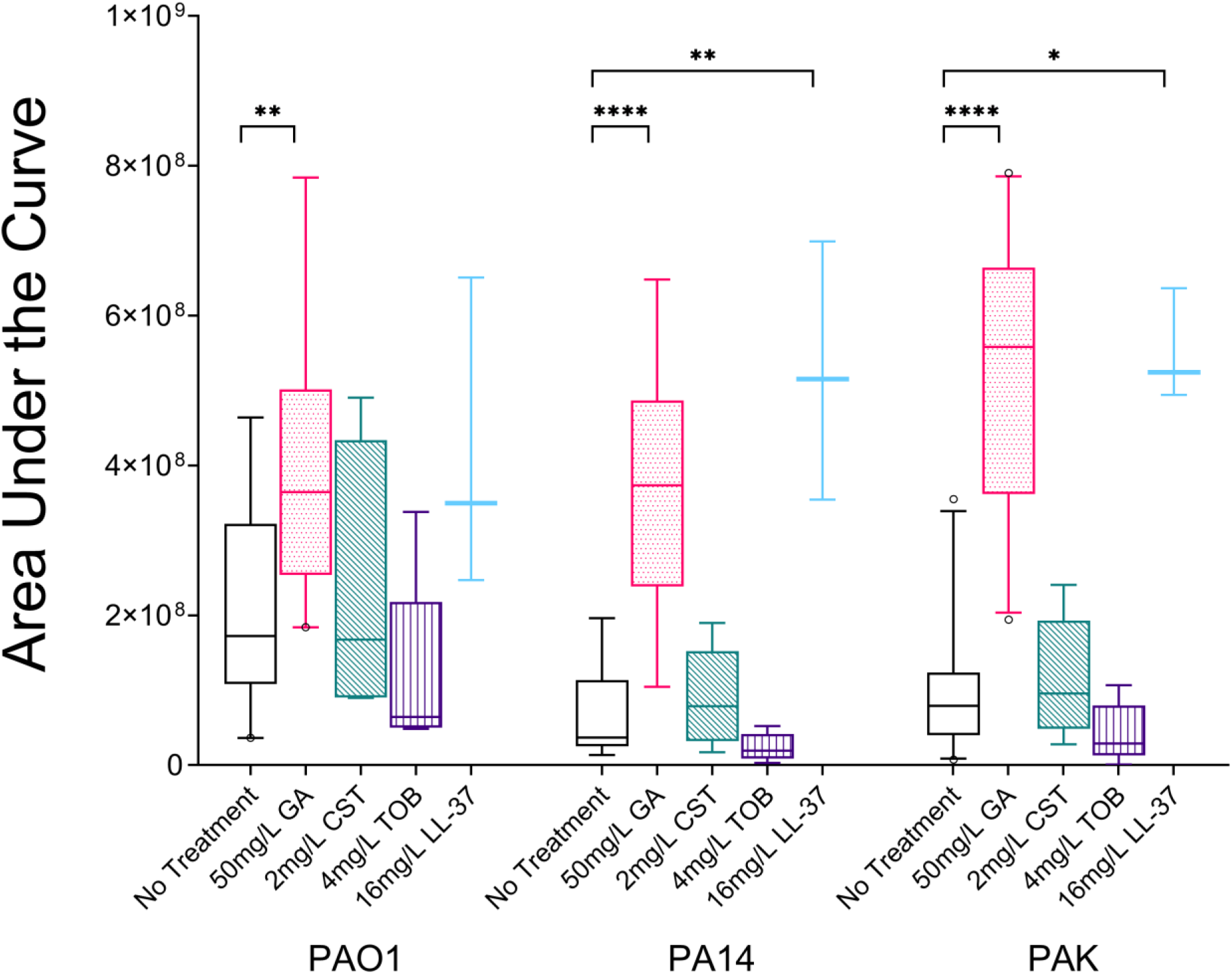
Comparison Areas Under the Curve of fluorescence of propidium iodide over 1hr of strains PAO1, PA14 and PAK resulting from exposure to 50mg/L GA, 2mg/L CST, 4mg/L TOB and 16mg/L LL-37 (Box and Whisker plot, line at median with 5-95 percentiles. At least three biological replicates of each treatment were performed for each strain). GA caused a significant increase in bacterial cell permeabilisation of all three reference strains (p<0.01).

## DISCUSSION

We hypothesised that the peptide drug, glatiramer acetate, would be an antibiotic resistance breaker in *P. aeruginosa*, when combined with the aminoglycoside, tobramycin. Here, we provide data to support this hypothesis and demonstrate the drug’s mechanism of action, using a single concentration of GA which has previously been shown to be optimally effective against *P. aeruginosa* (15). Initial tests with a small number of reference strains revealed that GA potentiated the activity of tobramycin as seen by bacterial growth, viability and MICs of these strains. Expanding these tests to clinical *P. aeruginosa* from people with cystic fibrosis confirmed these observations and resulted in significant reduction of TOB MICs; in the majority of TOB-resistant strains, the presence of GA rendered them TOB-sensitive. These effects likely relate to glatiramer acetate’s interaction with and disruption of the cell envelope of *P. aeruginosa* strains causing damage to the outer membrane, depolarisation of the cytoplasmic membrane and permeabilisation of the bacteria.

We employed a wide variety of techniques with both reference and clinical strains to fully understand the impact of GA. Using colony counts to calculate precise, granular minimum inhibitory concentrations we observed that glatiramer acetate was able to lower inhibitory concentrations required to reduce bacterial populations by 50% (MIC_50_) and 90% (MIC_90_). Glatiramer acetate synergised with tobramycin in the treatment of the 3 reference strains; the reduction in viable bacteria resulting from co-treatment was greater than would be expected, based on the individual treatments.

The ability to enhance or recover the activity of existing antibiotics is an extremely attractive property in a drug. An agent which is also already in clinical use and has moderate antibacterial activity is potentially desirable as a potential antibiotic adjuvant. It is crucial that studies such as this include clinical isolates, which are known to differ substantially from reference strains, at both genetic and phenotypic levels.

Encouragingly, GA/TOB synergy was also seen in the majority of strains in clinical *P. aeruginosa* from people with cystic fibrosis tested. This was the case regardless of whether the individual strain had been designated sensitive or resistant to tobramycin by standard clinical laboratory disk-diffusion testing. The isolates with both the lowest and highest MIC values of tobramycin were more susceptible to the aminoglycoside when GA was also present. This was not the case for every clinical isolate tested and we found examples of clinical strains which were unaffected in their sensitivity to tobramycin. Interestingly, this was not related to the resistance profile of these strains; both tobramycin-sensitive and -resistant *P. aeruginosa* isolates were among those with MICs minimally/not altered by co-treatment. Future work will focus on what these strains do have in common which may allow them to resist the potentiation of tobramycin by glatiramer acetate. Our data would appear to indicate that the two agents have distinct mechanisms of action given their ability to synergise, the lack of correlation between synergy and susceptibility to tobramycin and contrasting results from the two drugs in membrane perturbation assays.

Intrinsic antibiotic resistance in *P. aeruginosa* is closely related to the bacteria’s ability to exclude compounds from its cytoplasm, either via physical barrier (membrane impermeability) or toxin removal (efflux pumps). *P. aeruginosa*’s membrane is less permeable than that of other Gram-negative pathogens e.g. *Escherichia coli* (38–40). Indeed, in *P. aeruginosa* isolates from cystic fibrosis patients, “impermeability resistance” has been found to be the most common form of resistance to tobramycin (41,42). *P. aeruginosa* also encodes efflux pumps, such as MexXY, on its genome which can remove antibiotics, including tobramycin, from the cell interior (42–44).

In common with many other antimicrobial peptides, we found that glatiramer acetate disrupts the outer bacterial membrane, depolarises the cytoplasmic membrane and, ultimately, permeabilises the *P. aeruginosa* cell. The results reported here show that GA is as effective or more effective at causing membrane perturbations, in *P. aeruginosa* reference strains, than peptides colistin and LL-37. Among the agents tested, GA was the only treatment capable of causing significant changes to both membranes, which would allow entry to the cytoplasm, in all reference strains. Aminoglycosides have intracellular targets (ribosomal binding sites), where they can prevent protein production and cell replication (45). The activity of GA in disrupting the bacterial membrane may allow tobramycin access to these targets within the bacteria, by breaching the normally impermeable cell wall and/or overwhelming the activity of efflux pumps, preventing the cell from removing the antibiotic as quickly as it accumulates.

*P. aeruginosa* impermeability is aided by lipopolysaccharides (LPS) on the bacterial cell surface. *P. aeruginosa* can use divalent cations (e.g. Mg^2+^ and Ca^2+^) from their environment to bind together and stabilise the LPS chains forming an additional barrier to the bacterial membrane (46). In the absence of environmental cations or in response to certain stimuli, *P. aeruginosa* can also modify LPS chains, for example by addition of 4-amino-4-deoxy-L-arabinose (L-Ara4N), thereby changing the overall charge of the bacterial cell and making it more difficult for negatively charged antimicrobials (including tobramycin) to bind and enter the cell (47–49). Polycationic antimicrobials, such as glatiramer acetate, can displace the divalent cations which normally bind and stabilise LPS-LPS interactions and expose more of the bacterial membrane to external pressures/agents (50,51). Indeed, in the case of colistin, this targeting of LPS continues beyond the outer membrane and into the cytoplasmic membrane (33). The significance of divalent cations and LPS and their interactions has been known for over 50 years; the ability to potentiate antibiotics through cation and/or LPS removal and weakening of the defence they provide was demonstrated with the chelator, EDTA (52–56). This has made the desire to discover other, clinically safe “permeabilisers” which potentiate antibiotics a long standing one. We have demonstrated here glatiramer acetate’s ability to both permeabilise and potentiate *P. aeruginosa* to antibiotic treatment. Work is ongoing to investigate the specific interactions between GA and LPS structures and elucidate if GA potentiation works via a similar LPS interactions.

One of the leading drivers of antibiotic resistance and the reduction in effectiveness of existing antibiotics is the lack of development of new antibiotics. The crisis in the development of antibiotic resistance may also have been exacerbated by the COVID-19 pandemic, with secondary bacterial infections and poor antibiotic stewardship resulting in surges in antibiotic prescription (57,58). In the face of the dearth of investment in new antibiotics coming to market, alternative strategies are desperately needed (10,29). While there are important roles to be played by better testing, better antibiotic stewardship and novel strategies (bacteriophage, etc), these tactics do not directly address the effectiveness of currently available antibiotics. Resurrecting and/or improving the efficacy of clinically valuable antibiotics would be of great benefit, especially for an antibiotic as important as tobramycin. However, close attention will need to be paid to route of delivery of any novel treatment or adjuvant. People with cystic fibrosis already face a significant treatment burden (median 10 current treatments) and nebulised antibiotics are considered among the most burdensome of those treatments (59). Therefore, the most direct route to the CF clinic for an antibiotic adjuvant may be co-nebulisation with a partner antibiotic.

Antibiotic-resistant organisms being rendered ‘sensitive’ due the combination of therapies has obvious benefits to infected patients. Clinical benefits may include eradicating infections more quickly than with monotherapy (of both resistant and sensitive strains) and preventing chronic infections from becoming established. Ancillary benefits could accrue if potential antibiotic side-effects were reduced with shorter treatment courses or lower doses. Tobramycin has been associated with both ototoxicity and nephrotoxicity in cystic fibrosis, in a population with an already very high treatment burden of antibiotics (59–61). Whether or not the combined use of an antibiotic and a co-therapy such as GA would also have beneficial impact on resistance patterns emerging with time requires further study. While much attention is, naturally, focused the treatment of resistant bacteria, and we report MIC reductions and synergy with TOB-resistant *P. aeruginosa* here, our results also show that glatiramer acetate has similar synergistic and MIC-reducing capabilities with TOB-sensitive strains. This may also be clinically important due the heterogenous nature of the lung and the wide variation seen in nebulised tobramycin delivered to different regions (62,63). Reducing the concentration required to be effective against all *P. aeruginosa* types could be of benefit in the face of these concentration gradients within sputum, eradicating new infections and may aid in preventing the future development of resistant bacteria.

While further work is also required to ensure the effectiveness of GA in the complex environment of CF sputum and to formulate delivery to people with cystic fibrosis, the confluence of antimicrobial activity, tobramycin synergy and current clinical use makes glatiramer acetate a promising candidate for repurposing as an antibiotic adjuvant.

## Notes

### Competing Interest Statement

JH is an employee of Cycle Pharmaceuticals who provided Glatiramer Acetate as an in-kind contribution to the CF Trust grant for this work. GA is under license at the University of Aarhus, at which TVJ is employed.

