## Supplementary Figure 1 for "Synergistic Activity of Repurposed Peptide Drug Glatiramer Acetate with Tobramycin Against Cystic Fibrosis *Pseudomonas aeruginosa*"

Figure S1. Semilog plots of overnight growth of *P. aeruginosa* PA14 and PAK at 1mg/L TOB, with and without GA (natural logs of OD_600_, median and 95% confidence intervals).

Figure S1.

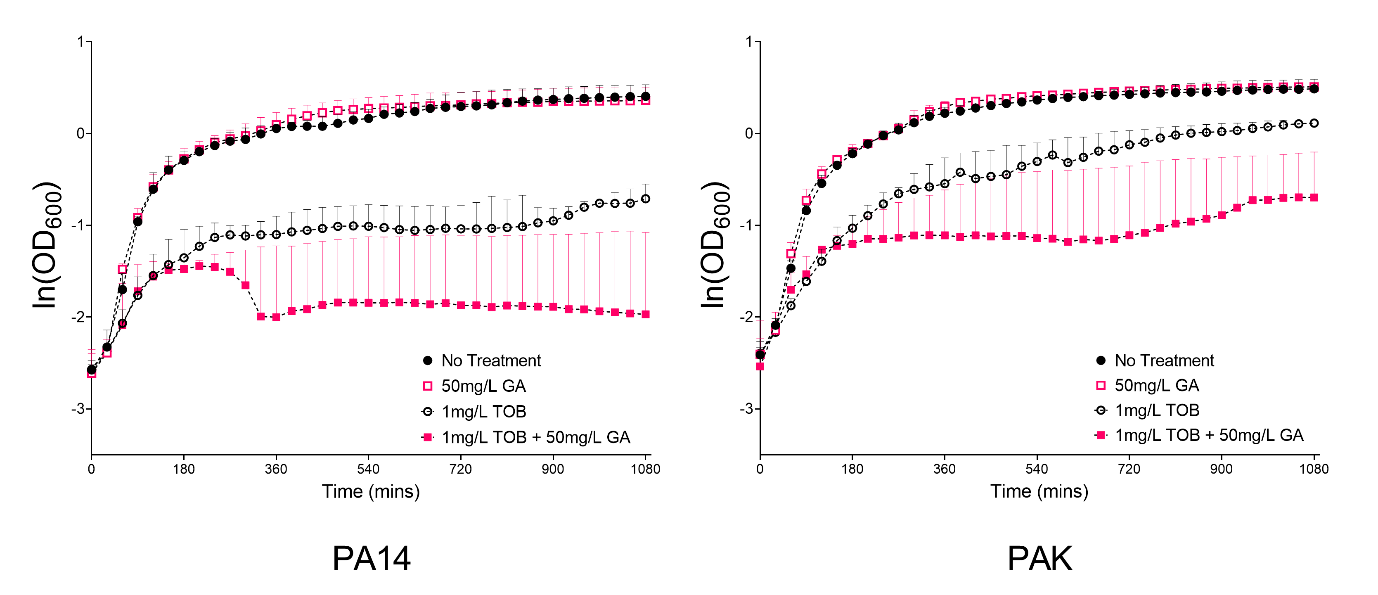
