## Supplementary Figure 2 for "Synergistic Activity of Repurposed Peptide Drug Glatiramer Acetate with Tobramycin Against Cystic Fibrosis *Pseudomonas aeruginosa*"

Figure S2. Inhibition Curves of *P. aeruginosa* strains PAO1, PA14 and PAK using Nonlinear Fit ([inhibitor] vs. response (three parameters)). Points show percent inhibition (of untreated control) (median and 95% confidence intervals). Tobramycin concentrations required to inhibit 50% (MIC_50_) and 90% (MIC_90_) of viable bacteria were interpolated from the curves generated for TOB-only and GA/TOB. Arrows indicate changes in the points where inhibition curves cross 50% and 90% when GA is present.

Figure S2.

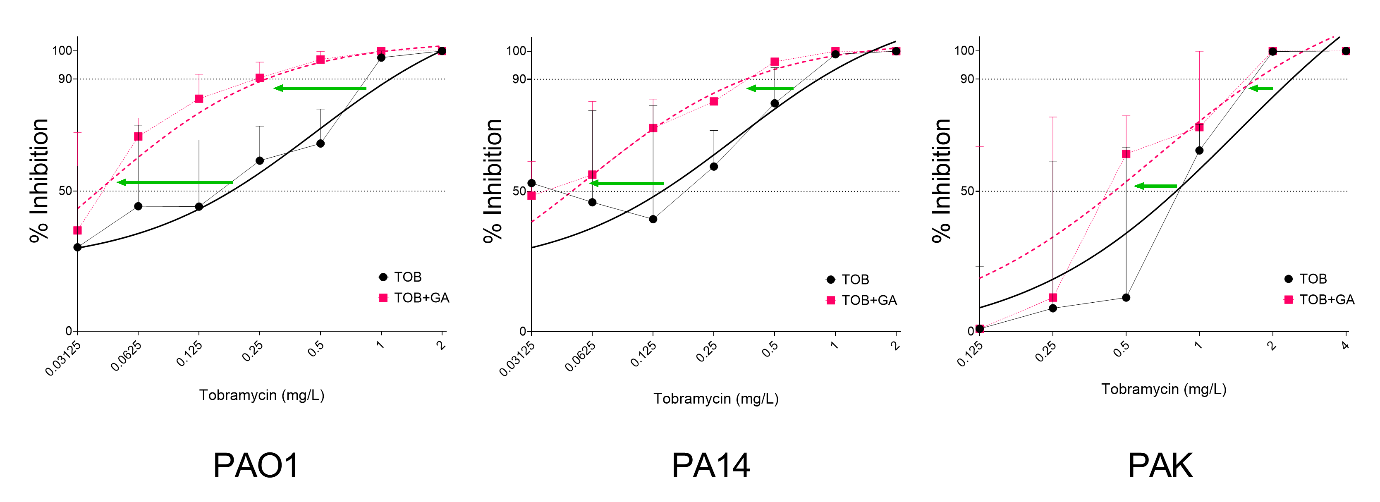
